# A Compact Two-Photon Module for Simultaneous Dual-Depth, Multi-Region *in vivo* Brain Imaging

**DOI:** 10.1101/2025.07.02.662755

**Authors:** Yaoguang Zhao, Xinyang Gu, Cihang Kong, Shoupei Liu, Yanfeng Zhu, Jixiong Su, Jingchuan Wu, Fang Xu, Liang Chen, Ying Mao, Bo Li

## Abstract

Understanding how distributed brain regions interact across cortical depths is crucial for uncovering the neural basis of behavior and cognition. Two-photon microscopy (2PM) excels at imaging the superficial layer, while three-photon microscopy (3PM) penetrates deep tissues but lacks sufficient speed for capturing fast superficial dynamics. Here, we present a compact two-photon imaging module (2PIM) that seamlessly integrates with existing 3PM to achieve simultaneous, high-speed imaging of superficial and deep brain regions, while preserving the deep tissue penetration of 3PM. This dual-modality system allows for remote and independent control of imaging planes in both axial and lateral dimensions. Leveraging this technology, we discovered context-dependent modulation of hippocampal CA1 and visual cortex responses to identical tactile stimuli, layer-specific temporal dynamics in sensory cortex during air puff stimulation, and depth-divergent neuronal activity patterns in motor cortex across acute-to-chronic pain. These results highlight the system’s potential for dissecting the spatiotemporal coordination of distributed brain circuits.

## 1. Introduction

Complex behaviors and cognitive states arise from the coordinated activity of spatially distributed neuronal populations across multiple regions and depths^1–5^. These interactions span diverse spatiotemporal scales from local microcircuit computations to long-range communication across cortical and subcortical networks^6–12^. Capturing such dynamics *in vivo* requires imaging technologies that combine deep tissue penetration, cellular resolution, high imaging speed and flexible multi-site targeting ^13, 14^.

Two-photon microscopy (2PM) is well-suited for high-speed, cellular-resolution imaging of superficial cortical layers but suffers from limited penetration depth^15–20^. In contrast, three-photon microscopy (3PM) enables imaging beyond 1 mm depth with reduced scattering and background fluorescence, making it ideal for subcortical access^21–25^. However, the intrinsically low repetition rate of 3PM excitation laser constrains multi-plane acquisition speed, particularly limiting its performance for tracking fast activity in superficial areas. Consequently, neither modality alone is sufficient for simultaneous, rapid imaging across deep and superficial regions—a fundamental limitation in systems neuroscience.

Hybrid approaches that combine 2PM and 3PM have attempted to overcome this limitation but typically permit only axial (depth-wise) displacement of imaging planes^25–29^. In realistic experimental settings, however, regions of interest often differ not only in depth but also in lateral location, complicating simultaneous imaging. Moreover, these systems often require extensive optical modifications or rely on custom-built microscope platforms, limiting their adaptability and scalability. Alternative methods, such as gradient refractive index (GRIN) lens implantation, allow access to multiple depths but involve invasive surgery, fixed focal paths, and restricted lateral coverage ^13, 14, 30^. Thus, a dual-region imaging solution that is modular, minimally invasive, and capable of remote, independent axial and lateral control—without compromising speed or compatibility— is urgently needed.

To this end, we developed a compact, modular two-photon imaging module (2PIM) that seamlessly integrates with standard 3PM. This dual-modality system enables simultaneous, high-speed imaging of both superficial and deep brain regions without sacrificing temporal resolution. Importantly, 2PIM provides independent axial and lateral control of the two-photon (2P) imaging planes, allowing flexible targeting of spatially disparate brain areas in real time. Leveraging this system, we observed context-dependent modulation of visual and hippocampal responses to identical tactile stimuli, revealed layer-specific sensory dynamics, and identified depth-resolved neuronal adaptations in chronic pain models. These findings establish 2PIM as a versatile platform for investigating the spatiotemporal organization of distributed brain circuits underlying complex behaviors and cognitive states.

## 2. Results

### 2.1 Modular Design of a Compact 2PIM for Multi-Region Imaging with 3PM

To develop a multi-area brain imaging module that can be rapidly integrated into existing systems, we designed a compact 2PIM. The 2PIM sequentially modulates the 2P laser beam via three key submodules: time-division multiplexing (TDM), remote focusing (RF), and remote lateral shift (RLS) (**Figure 1**a and **S1**). These components respectively manage temporal gating, axial focusing, and lateral positioning of the 2P beam. The modulated beam is then optically combined with the three-photon (3P) excitation path via a dichroic mirror before the galvanometer, enabling multi-area imaging.

**Figure 1.**
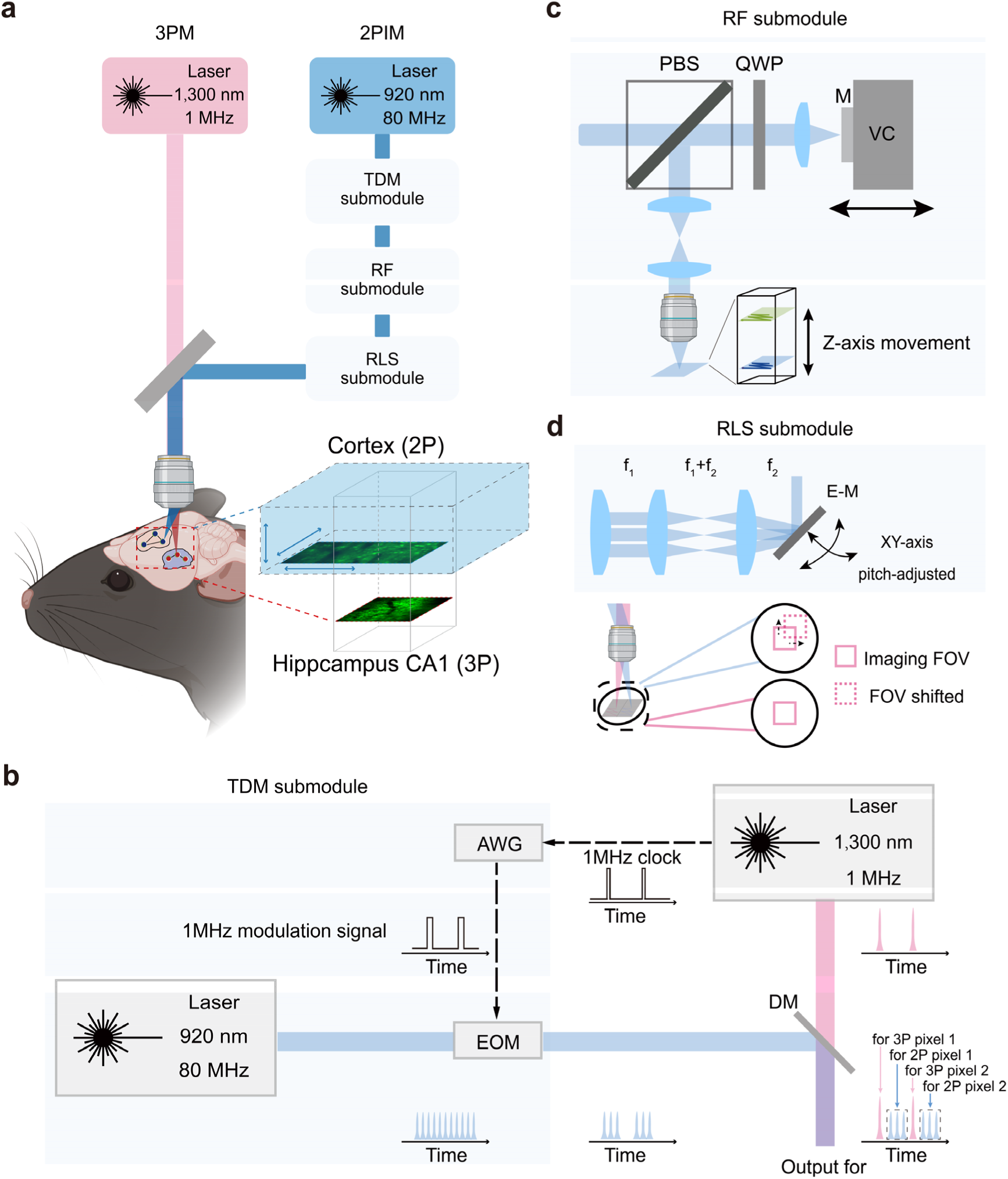
Schematics of the 2PIM and 3PM for multi-region imaging. a) Conceptual diagram of 3P imaging compatibility with the 2PIM. b) Schematic of the TDM submodule. AWG: arbitrary waveform generator; EOM: electro-optic modulator; DM: dichroic mirror. This submodule enables crosstalk-free 2P/3P simultaneous imaging through pulsed-laser modulation. c) Schematic of the RF submodule. PBS: polarization beam splitter; QWP: quarter-wave plate; M: mirror; VC: voice coil. This submodule modulates beam divergence to adjust the 2P focal plane along the Z-axis. d) Schematic of the RLS submodule. E-M: electrically controlled mirror. This submodule can adjust the E-M angle to shift the 2P imaging FOV laterally.

First, we developed the TDM submodule to enable synchronized dual-modality imaging (Figure 1b). This submodule inserts modulated 2P pulses into the temporal gaps between consecutive 3P pulses, allowing both 2P and 3P images to be sequentially acquired within the dwell time of a single pixel. This design offers two key advantages: (1) the acquisition intervals for modalities occur within ∼0.5 μs, enabling near-simultaneous sampling of distinct imaging planes, and (2) the 2P image is captured during the acquisition interval of the 3P image, thereby preserving the imaging speed and resolution of 3P imaging, which is especially crucial for low-repetition-rate 3P systems. Technically, the 1 MHz synchronization clock of the 3P laser serves as a timing trigger for an arbitrary waveform generator (AWG), which in turn modulates the 2P laser pulse train via an electro-optic modulator (EOM). By tuning the AWG’s delay and waveform shape, we achieved precise temporal alignment and control over the burst profile of the 2P pulses, thereby minimizing crosstalk between the two imaging modes (**Figure S2**).

Next, we designed a RF submodule that provides independent axial control of the 2P imaging plane without interfering with the 3P optical path (Figure 1c). This module employs a mirror positioned at the conjugate plane of the scan focus, which is actuated by a voice coil motor to adjust the axial (Z-axis) position of the 2P focal plane in real time. Because this configuration is optically decoupled from the 3P excitation path, it enables dynamic focal depth modulation of the 2P beam without perturbing the 3P imaging volume, a critical requirement for multi-region imaging. For example, during simultaneous imaging of the cortex and hippocampus, the 3P modality can target deep CA1 neurons, while the 2P plane can be independently tuned to superficial cortical layers. The RF submodule consists of a set of optical components, including polarizing beam splitters (PBS), quarter-wave plates (QWP), lenses (L), and a reflective mirror (M), mounted on a linear translation stage driven by a voice coil (VC) motor under software control. Within its operational axial range (ΔZ = 0-0.6 mm), we observed a consistent and previously undescribed change in the 2P field of view (FOV, both X and Y axes, **Figure S3**). Notably, this change modestly expands the 2P imaging area, albeit with a slight reduction in spatial sampling resolution.

Additionally, we developed an RLS submodule that enables independent and real-time precise control of the lateral (X-Y) position of the 2P imaging plane (Figure 1d). This submodule incorporates an electrically actuated mirror placed between the galvanometer scanner and the RF submodule, which is conjugate to the scanning focal plane. By fine-tuning the mirror angle, the 2P beam can be steered laterally without affecting the optical path or focal alignment of the 3P imaging system. This optical decoupling enables real-time lateral translation of the superficial imaging plane. Such flexibility is critical for simultaneous targeting of anatomically distinct regions that may differ not only in depth but also in lateral position. In practice, once the desired depth is established using the RF submodule, the RLS submodule enables seamless repositioning of the 2P plane to interrogate nearby structures without physically moving the sample or realigning the optical system. This capability significantly enhances spatial coverage and experimental throughput in multi-area recordings, particularly in awake animal paradigms where stability and speed are paramount.

Finally, by combining the 2P and 3P excitation paths through a dichroic mirror, along with the TDM, RF, and RLS submodules, we constructed a fully modular multi-area imaging system. This dual-modality platform enables simultaneous, high-speed acquisition from spatially distinct brain regions with independent control over both axial and lateral positioning of the 2P imaging plane.

### 2.2 Functional Validation of the Integrated 2PIM-3PM Platform for Multi -Region Brain Imaging

We integrated the 2PIM module and its laser path into a 3PM and first evaluated the performance of the TDM submodule. When only the 3P excitation was active (**Figure 2**a, top), distinct 3P laser pulses and their corresponding fluorescence signals were reliably detected in the 3P channel (Channel 1), with no observable signal in the 2P channel (Channel 2), confirming the absence of crosstalk. Conversely, when only the modulated 2P excitation was applied (Figure 2a, middle), characteristic 2P fluorescence signals were successfully recorded in the 2P channel. Importantly, under simultaneous 2P and 3P excitation, both signal streams were acquired concurrently on a unified timeline (Figure 2a, bottom). These results confirm that the TDM submodule effectively enables clean temporal separation and concurrent acquisition of 2P and 3P signals within the same imaging session.

**Figure 2.**
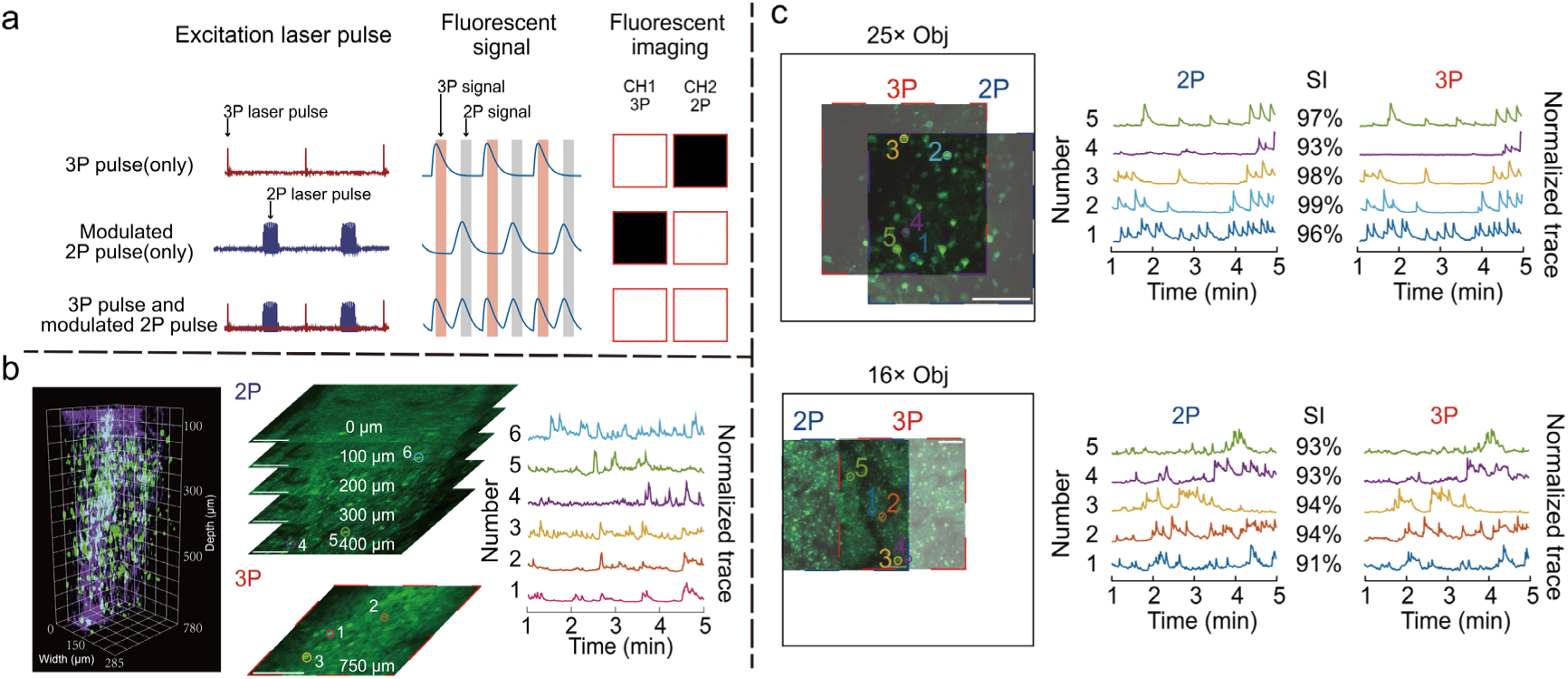
Functional validation of three core submodules: TDM, RF, and RLS. a) TDM performance evaluation. Top: 3P only; middle: modulated 2P only; bottom: simultaneous 2P/3P. Left: laser pulse signals (purple: 2P excitation at 920 nm; red: 3P excitation at 1,300 nm); center: fluorescent signals (orange: 3P window; grey: 2P window); right: fluorescent solution imaging (CH1: channel 1; CH2: channel 2). b) RF axial scanning capability. Left: 3P volumetric imaging of mouse somatosensory cortex (depth: 0-780 μm); center: 2P imaging at varying depths when 750 µm 3P focus fixed; right: extracted neuronal trace at corresponding axial planes. c) RLS lateral field adjustment. Top row: 25× objective; left: average-projection image; right: calcium traces; bottom row: 16× objective; SI: similar index between 2P and 3P. Scale bar in b and c: 100 μm.

We next evaluated the system’s ability to independently control the axial movement of the 2P imaging plane. Initially, we employed 3P imaging to capture 3D structural information from a 780 μm deep region of the mouse brain (Figure 2b left). After fixing the depth of the 3P imaging at 750 μm, we used the RF submodule of the 2PIM to adjust the depth of the 2P imaging plane, capturing data across five depths ranging from 0 to 400 μm (Figure 2b, middle). Robust calcium transients with high signal-to-noise ratios were consistently recorded at different depths (Figure 2b middle right), confirming that the system could independently control the vertical movement of the 2P imaging plane, while maintaining high-quality data acquisition at different depths.

We then assessed the system’s capability for independent lateral movement of the 2P imaging plane. In this experiment, the imaging depth of both the 2P and 3P modalities was maintained constant while using a 25× water-immersion objective (Olympus, NA = 1.05). Under these conditions, the 3P imaging FOV was 285 μm × 285 μm, while the 2P imaging FOV was identical but could be freely moved within the objective supported FOV (around 500 μm × 500 μm) (Figure 2c top and **Figure S4** top). Calcium activity traces extracted from the same neurons using both imaging modes were nearly identical. This strong agreement is primarily attributed to the near-simultaneous sampling of both modalities, further confirming that independent lateral movement of the 2P imaging plane does not interfere with TDM or RS performance. The range of 2P lateral translation is primarily determined by the objective’s maximum FOV. For example, when using a 16× objective (Nikon, NA = 0.8), the 2P imaging plane could move freely within a larger area (> 1,000 μm × 1,000 μm, Figure 2c bottom and Figure S4 bottom). If further displacement is needed, lower-magnification, large-FOV objectives can support millimeter-scale lateral repositioning, as demonstrated in prior mesoscopic imaging studies^31–33^.

### 2.3 Functional Validation of Dual-Regional Imaging in Cortex and CA1 Under Aversive Stimuli

To validate the system’s capability for multi-area brain imaging, we simultaneously recorded neuronal activity in the visual cortex (500 μm depth, 2P) and hippocampal CA1 (1,000 μm depth, 3P) during a combined visual and air puff stimulation paradigm (**Figure 3**a and 3b). High-fidelity calcium traces from four representative neurons were stably recorded over a continuous one-hour session, demonstrating the system’s capacity for reliable, depth-resolved signal acquisition. We further quantified calcium activity across all detected neurons by measuring photon counts, signal discriminability (*d’*), and relative fluorescence changes Δ*F/F* (Figure 3c). Notably, although photon counts were high in 2P imaging at 500 μm depth, Δ*F/F* values remained low due to elevated background fluorescence. To mitigate this, we assumed a spatially uniform background signal as a direct current (DC) component and generated background-subtracted (BS) 2P images (see Methods). We then compared three imaging conditions: conventional 2P, BS-2P, and 3P. Across all modalities, *d’* values exceeded or approximated 5, corresponding to calcium detection accuracy of ∼96%^34^.

**Figure 3.**
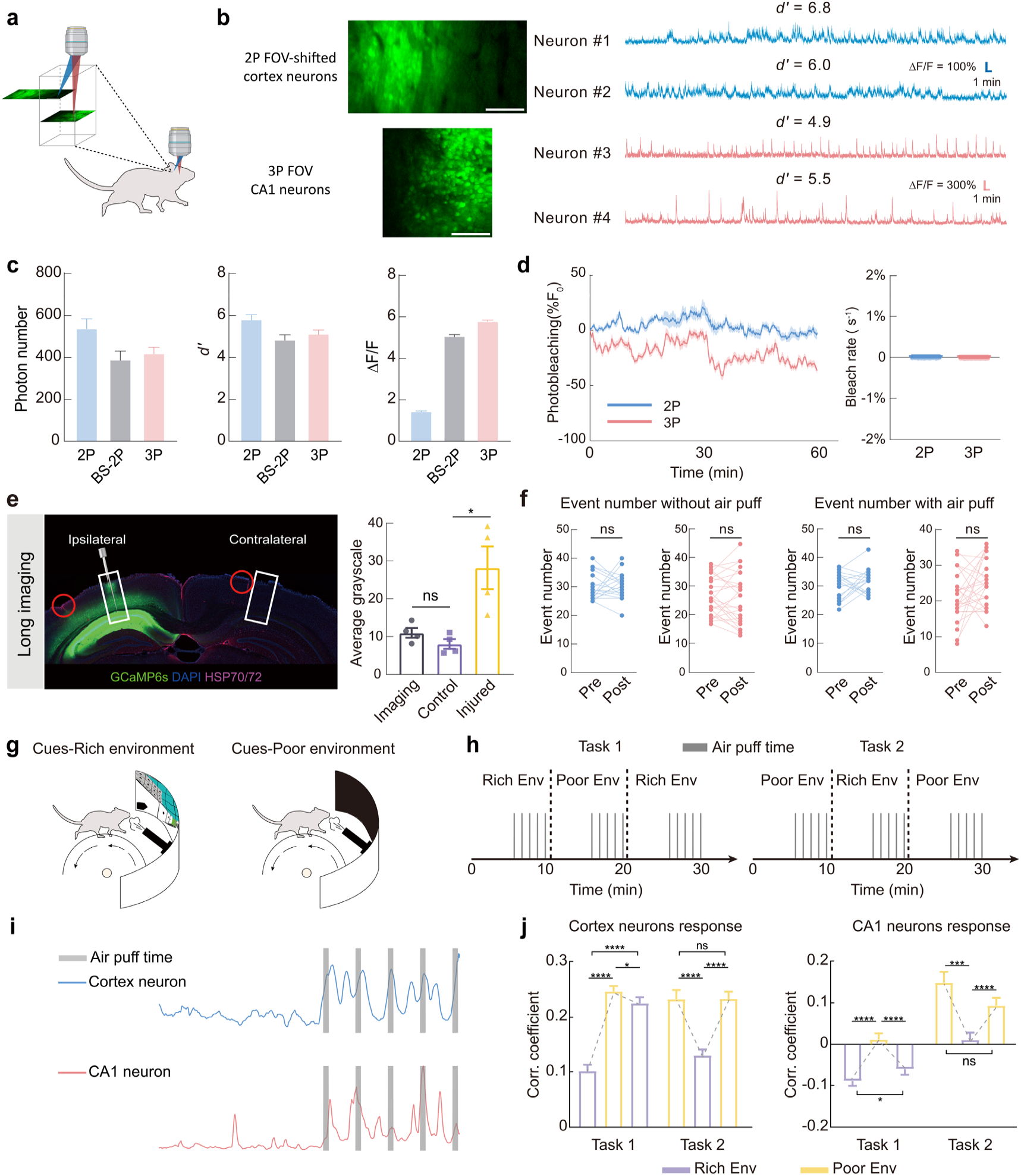
Validation and application of dual-region imaging in cortex and CA1. a) Imaging paradigm. b) Simultaneous 2P and 3P imaging demonstration. Left: representative images; right: calcium traces (Δ*F/F*) from representative cortical and CA1 neurons recorded over a 1-hour session, and corresponding *d’* values are indicated. c) Quantitative metrics of simultaneous 2P and 3P imaging. Left: photon counts for 2P, BS-2P, and 3P imaging; middle: *d’*; right: Δ*F/F*. d) Photobleaching analysis during 1-hour imaging. Left: bleaching curves for cortex (2P) and CA1 (3P), plotted as mean ± SEM; right: derived bleaching rates. e) Assessment of thermal damage using HSP expression. Grayscale intensity values were compared across the imaging region, contralateral (non-imaged) region, and injured (caused by surgery) control. Statistical method: unpaired two-tailed t-test, *n* = 4 FOVs, *P* (Imaging vs. Control) = 0.1589; *P* (Control vs. Injured) = 0.0132. f) Neuronal calcium event number before and after 1-hour imaging. Left: without air puff; right: with air puff. Statistical method: paired two-tailed t test, *n* (2P) = 18 neurons, *n* (3P) = 25 neurons, *P* = 0.8213, 0.5238, 0.3680, 0.1791. g) Schematic of the mouse virtual reality (VR) behavioral paradigm and dual-region imaging setup. h) Task design. Cues-Rich environment (Rich Env) and Cues-Poor environment (Poor Env) were alternated every 10 minutes. Task1: Rich-Poor-Rich. Task2: Poor-Rich-Poor. Air puff stimuli were delivered during the final 5 minute of each environment. i) Representative neuronal responses to air puff stimuli. j) Response intensity analysis based on Pearson correlation. Left: response intensity of cortical neurons under tasks; right: corresponding data for CA1 neurons. Statistical methods: paired two-tailed t-test, *n* (2P) = 77 neurons (2 mice), *P* (Poor 1 vs. Rich 2) = 0.0323, *P* (Poor 2 vs. Poor 3) = 0.9653, \*\*\*\**P* < 0.0001; *n* (3P) = 110 neurons (2 mice), *P* (Rich 1 vs. Rich 2) = 0.0308, *P* (Poor 2 vs. Rich 3) = 0.0002, *P* (Poor 2 vs. Poor 3) = 0.0742, \*\*\*\**P* < 0.0001. Statistical result shown in e and d is mean ± SEM, and in c and j are mean + SEM.

We evaluated the photostability of the system by quantifying photobleaching kinetics across all recorded neurons during a continuous one-hour imaging session (Figure 3d; see Methods). The average photobleaching rates for both modalities were negligible, measured at 0.00% s^-1^ for 2P imaging and 0.01% s^-1^ for 3P imaging, indicating minimal photobleaching under the applied conditions.

To assess potential phototoxicity, we examined heat shock protein (HSP) expression as a biomarker for thermal damage. Comparative analysis between imaged and contralateral non-imaged regions revealed no significant differences in HSP expression (Figure 3e), suggesting that prolonged imaging does not induce detectable thermal damage. Despite the absence of histological damage, we further investigated whether the imaging procedure perturbs neuronal function. In the absence of stimulation (without air puff condition), neuronal activity remained stable over time, with no significant difference in the number of calcium events between the first and last 1,000 imaging frames (Figure 3f, left). Quantitatively, visual cortex neurons (imaged with 2P) exhibited an average of ∼30 events per session, and CA1 neurons (3P) showed a comparable event count. Similarly, during air puff stimulation, neuronal activity levels remained consistent across time (Figure 3f right), with visual cortex and CA1 neurons averaging ∼30 and ∼20 events per session, respectively.

We further leveraged our dual-modality imaging system to investigate how environmental visual context modulates neural responses to aversive stimuli across multiple brain regions. Previous studies have demonstrated that hippocampal CA1 pyramidal neurons exhibit significant spatial remapping following contextual fear conditioning, especially when aversive stimuli (e.g., air puffs, foot shocks, or predator odors) are introduced into familiar spatial representations^35–38^. Moreover, CA1 neurons are known to encode aversive stimuli independently of spatial location, with their response intensity modulated by tactile contexts^39^. Given the critical role of visual inputs in environmental perception alongside tactile cues, we utilized our system to simultaneously record neuronal activities from the visual cortex and hippocampal CA1 while exposing mice to identical air puff stimuli across visually distinct environments (Figure 3g). In this paradigm (Figure 3h), head-fixed mice freely ran on a treadmill during imaging sessions. The surrounding visual environment—provided via a frontal screen—was systematically altered every 10 minutes, with each 10 minutes interval defined as a “session”. Air puff stimuli were delivered five times during the final 5 minutes of each session. Every task contains 3 sessions, and two task conditions were employed: Task 1 followed a Cues-Rich environment → Cues-Poor environment → Cues-Rich environment order, while Task 2 followed a Cues-Poor environment → Cues-Rich environment → Cues-Poor environment order.

Across both visual and hippocampal regions, air puff stimuli elicited robust calcium responses (Figure 3i). Notably, the response amplitudes exhibited a clear inverse correlation with the richness of visual cues: weaker responses were observed in Cues-Rich conditions, while Cues-Poor conditions led to significantly stronger activations (Figure 3j). This effect persisted regardless of task sequence, indicating that neuronal responses were modulated by environmental context rather than exposure order. Contextual modulation was more pronounced in the visual cortex: visual dominance in Cues-Rich settings appeared to suppress somatosensory-driven responses, whereas visual deprivation in Cues-Poor settings enhanced neural sensitivity to tactile stimuli. CA1 showed a similar but weaker modulation pattern, suggesting a potential role in integrating cross-modal information to facilitate spatial remapping.

### 2.4 2PIM Dissects Layer-Resolved Temporal Dynamics in Sensory Cortex Under Air Puff Stimulation

Emotional states significantly influence human health and disease outcomes. However, how the brain converts experiences into emotions remains unclear^40^. Understanding the neural basis of emotional responses (e.g., positive vs. negative feelings) is a core challenge in neuroscience. This complexity arises because emotions depend on two key factors: (1) the brain’s encoding of emotionally relevant information, and (2) modulation by sensory inputs, internal bodily sensations, and neurochemical systems^41^. Consequently, deciphering the neurocircuitry mechanisms governing negative valence bears fundamental significance for understanding behavioral pathologies and mental disorders.

Within cortical microcircuits, Layer V pyramidal neurons serve as the primary output projection system, whose synchronized firing drives long-range axonal signaling to thalamus-subcortical targets that orchestrate arousal, attentional allocation, and sensorimotor integration^42^. In contrast, Layer II-III neurons predominantly engage in local microcircuit computations, with their coordinated activity enabling sensory feature extraction and cross-regional synchronization^43, 44^. The neural encoding of air puff stimulation constitutes a complex spatiotemporal pattern transformation process from peripheral receptors to the cortex^45, 46^. The primary somatosensory cortex (S1) utilizes intricate intracortical microcircuits to extract multidimensional features of air puff stimuli—including spatial location, intensity, frequency, and directional movement—which are represented through distributed activity patterns of population neurons. Consequently, investigating information processing and collaborative coding mechanisms in higher neural centers—using air puff-evoked transient negative valence responses as a model system—provides a crucial window into fundamental principles of emotional information processing.

To dissociate layer-specific neuronal roles in emotional processing, we designed a behavioral paradigm (**Figure 4**a): head-fixed mice were allowed to run freely on a wheel while receiving 10s air puff stimuli to the whisker pad and neuronal calcium activity was simultaneously recorded in superficial (∼300 μm, layer II-III, 2P) and deep (∼700 μm, layer V, 3P) somatosensory cortex. We found both layers exhibited response heterogeneity: subsets of neurons were positively responsive, non-responsive, or negatively responsive to air puff (Figure 4b).

**Figure 4.**
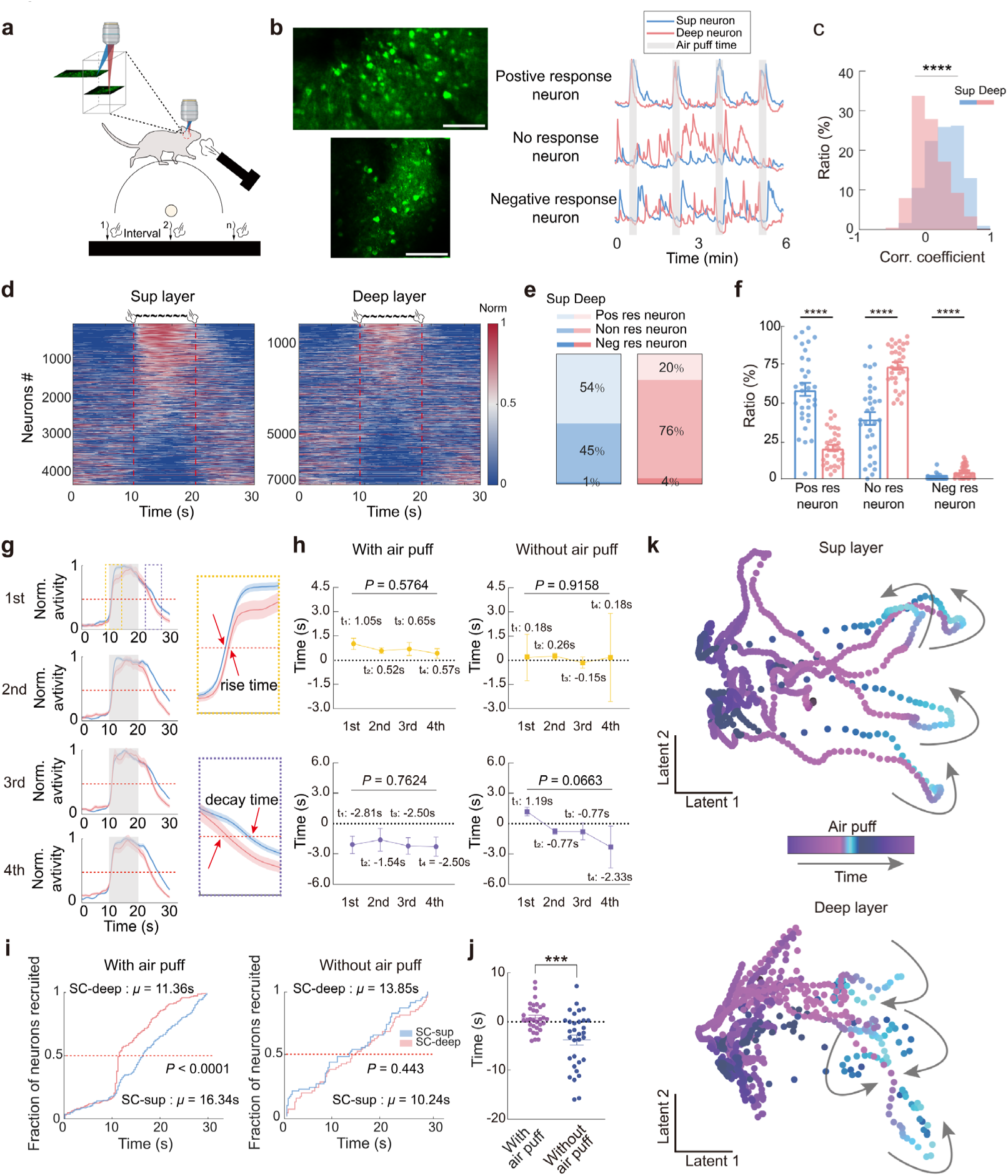
Functional differences in sup and deep neuronal activities in the somatosensory cortex induced by air puff stimulation using 2PIM. a) Behavioral paradigm for imaging. b) Imaging demonstration and neuronal activity. Top left: superficial layer (abbreviated as “Sup” in figure hereafter); bottom-left: deep layer; right: neurons with different responses to air puff stimulation (pos res: positive response; neg res: negative response; non res: non response). Scale bar: 100 μm. c) Distribution of Pearson correlation coefficients between neuronal activity and air puff stimulation in superficial (blue) and deep (orange) layers. Statistical method: Kolmogorov-Smirnov test, *n* (2P) = 1077 neurons (4 mice), *n* (3P) = 1764 neurons (4 mice), *P* <0.0001. d) Heatmaps of population activity over 30s before and after air puff stimulation. Neurons were sorted by correlation. Left: superficial layer; right: deep layer. e) Proportions of different neuron types in superficial and deep layers. f) Quantification of neuron type proportions. Statistical method: two-tailed Mann-Whitney U test, *n* = 32 FOVs (4 mice), \*\*\*\**P* < 0.0001. g) Average neuronal traces within 30s around stimulation. Left: traces of the 1st–4th stimuli (top to bottom), normalized to [0,1]; right: an enlarged view of the first stimulus (yellow box: rising phase; purple box: decaying phase). h) Quantification of temporal differences in rising and decaying phases between superficial and deep layers. Top left: rising; bottom left: decaying; right: the results of without air puff as control. Statistical method: Kruskal-Wallis test, *n* = 32 FOVs (4 mice), *P* values as shown. i) Cumulative distribution curves of active neurons. Left: with air puff stimulation; right: without air puff stimulation as control. Statistical method: Kolmogorov-Smirnov test, *P* values as shown. j) Quantification of differences in the time to reach 50% activation between superficial and deep neurons. Left: with air puff; right: without air puff. Statistical method: unpaired two-tailed test, *n* = 32 FOVs (4 mice), *P* = 0.0002. k) High-dimensional space characteristics of positive response neurons. Top: PCA dimensionality reduction of superficial neurons over time (blue: air puff periods, 4 stimuli); bottom: the deep layer. Statistical result shown in f and g is mean ± SEM.

In order to test if this heterogeneity represents a stable phenomenon rather than the random firing of individual neurons coinciding with an air puff. We calculated Pearson correlation coefficients between neuronal activity in superficial/deep layers and air puff stimuli. The results showed significantly different correlation coefficient distributions between layers (Figure 4c), suggesting functional specialization across sensory cortex depths during air puff responses. To visually characterize this functional heterogeneity, we generated normalized activity heatmaps of single-trial neuronal responses in deep and superficial layers (Figure 4d) and performed correlation-based neuronal classification (Figure 4e). This analysis revealed striking layer-specific differences in the proportions of positive response (54% in superficial vs. 20% in deep), no response (45% in superficial vs. 76% in deep), and negative response neurons (1% in superficial vs. 4% in deep). In order to verify the universality of layer-specificity, we calculated the 32 FOVs in different trials for 4 mice, and single-FOV unit analyses (Figure 4f) yielded consistent results, confirming the reliability of inter-layer differences. Analysis of fluorescent intensity dynamics across the three neuronal classes during distinct phases of air puff stimulation further revealed divergent threat response patterns (**Figure S5**a-b and c).

We found that superficial positive response neurons exhibited faster rising but slower decay than deep neurons upon stimulation (Figure 4g). Quantifying time differences to reach 50% intensity within the same FOV (Figure 4h left) revealed that during the first stimulus, superficial layer activated 1.05 ± 0.36 s (mean ± SEM, likewise hereafter) earlier and decayed 2.81 ± 1.2 s slower than deep layer. With repeated stimuli, rise time delays stabilized at 0.57 ± 0.36 s and decay delays at -2.50 ± 0.96 s. Control analysis of inter-stimulus intervals (Figure 4h right) showed no significant differences in rise time (0.17 ± 1.44s vs. 0.18 ± 2.7 s) or decay time (1.2 ± 0.43s vs. -2.3 ± 2.1 s) between epochs, but their coefficient of variation was significantly higher than during stimulation (Figure S5d), confirming temporal stability of air puff-induced effects. When disregarding response specificity and conducting analysis at the aggregate response level, we validated these layer-specific temporal dynamics (Figure 4i): the activation time of 50% of deep neurons during air puff stimulation significantly differed from superficial neurons (deep is ∼5s faster), whereas control distribution curves showed no differences. Quantification of mean differences across 4 mice, including 32 different FOVs, (Figure 4j) revealed a significant distinction between stimulus-period time differences (0.85 ± 0.51 s) and non-stimulus-period differences (-3.70 ± 1.0 s), further confirming the specificity of stimulus-related responses. These findings suggest that superficial neurons mediate threat stress regulation through rapid activation and prolonged signaling, while deep neurons prioritize rapid signal integration and baseline recovery after threat processing.

To explore high-dimensional dynamics of neuronal population activity, we performed principal component analysis (PCA) for dimensionality reduction. Results showed that the population activity of positively responsive neurons exhibited transition trajectories between two subspaces in high-dimensional space (Figure 4k), with more pronounced migration trajectories in superficial layer, likely associated with their functional complexity. Non-responsive and negatively responsive neurons showed no significant state-dependent differences (Figure S5e), revealing feature segregation of functionally distinct neuron types in high-dimensional space.

In summary, simultaneous dual-depth imaging of superficial and deep neurons in the somatosensory cortex reveals a layer-stratified computational architecture for processing negative valence. Superficial layer exhibits high-ratio recruitment and prolonged activity maintenance, suggesting their role in sustained threat representation and cortico-cortical integration of emotional context. In contrast, deep layer demonstrates rapid signal onset and accelerated decay, indicating their specialization in transient threat detection and fast feedback to subcortical effectors for immediate behavioral adaptation. This functional segregation establishes a cortical microcircuit mechanism where superficial layer sustains affective states for cognitive evaluation, while deep layer prioritizes rapid disengagement to restore homeostasis. Crucially, the divergent high-dimensional state trajectories—particularly the pronounced migration in superficial layer—reflect their dynamic reconfiguration during threat processing, aligning with their proposed role in integrating multisensory inputs for flexible emotion regulation.

### 2.5 2PIM Reveals Layer-Specific Motor Cortex in Response to Chronic Pain

Traditional pain theory divides nociception into three dimensions: sensory, affective, and cognitive^47, 48^. The involvement of the primary motor cortex (M1) reveals the core role of the “motor dimension”^49^. As a hub integrating motor and nociceptive processing, M1 competitively inhibits ascending pain signals by activating non-nociceptive sensory inputs (e.g., touch), participates in placebo effects and endogenous opioid release, thereby integrating the motor system into the core of the “pain matrix”.

Long-standing research indicates that chronic pain arises not only from sensory pathway dysfunction but also from dysfunctional interaction between sensory and motor systems^50^. M1 circuits concurrently regulate both sensory and affective dimensions. Deciphering the neural encoding of M1 under chronic negative stimuli in pain models provides a solid foundation for addressing motor-pain comorbidities^51^.

Previous studies have revealed complex neural encoding mechanisms of negative valence in sensory cortices. This study aims to investigate how the motor cortex responds to sustained negative valence induced by prolonged nociceptive stimulation in rodent pain models. We employed a spared nerve injury (SNI) model (validity confirmed in **Figure S6a**) to decode functional dynamics across cortical depths of the M1. As shown in **Figure 5a**, C57BL/6JNifdc wild-type mice received intraparenchymal injections of adeno-associated virus AAV2-CaMKII-GCaMP6s 30 days before SNI surgery to label neurons with the fluorescent calcium indicator GCaMP6s; craniotomy was performed 20 days before modeling. *In vivo* longitudinal imaging was conducted on pre-modeling (D0) and post-modeling days 1, 7 and 14: head-fixed mice were allowed to run freely on a wheel, while simultaneously recording locomotion distance, speed, and calcium activity in superficial (∼200 μm, layer II) and deep (∼600 μm, layer Ⅴ) M1 cortex to acquire high-quality neuronal activity signals for subsequent analysis (Figure 5b).

**Figure 5.**
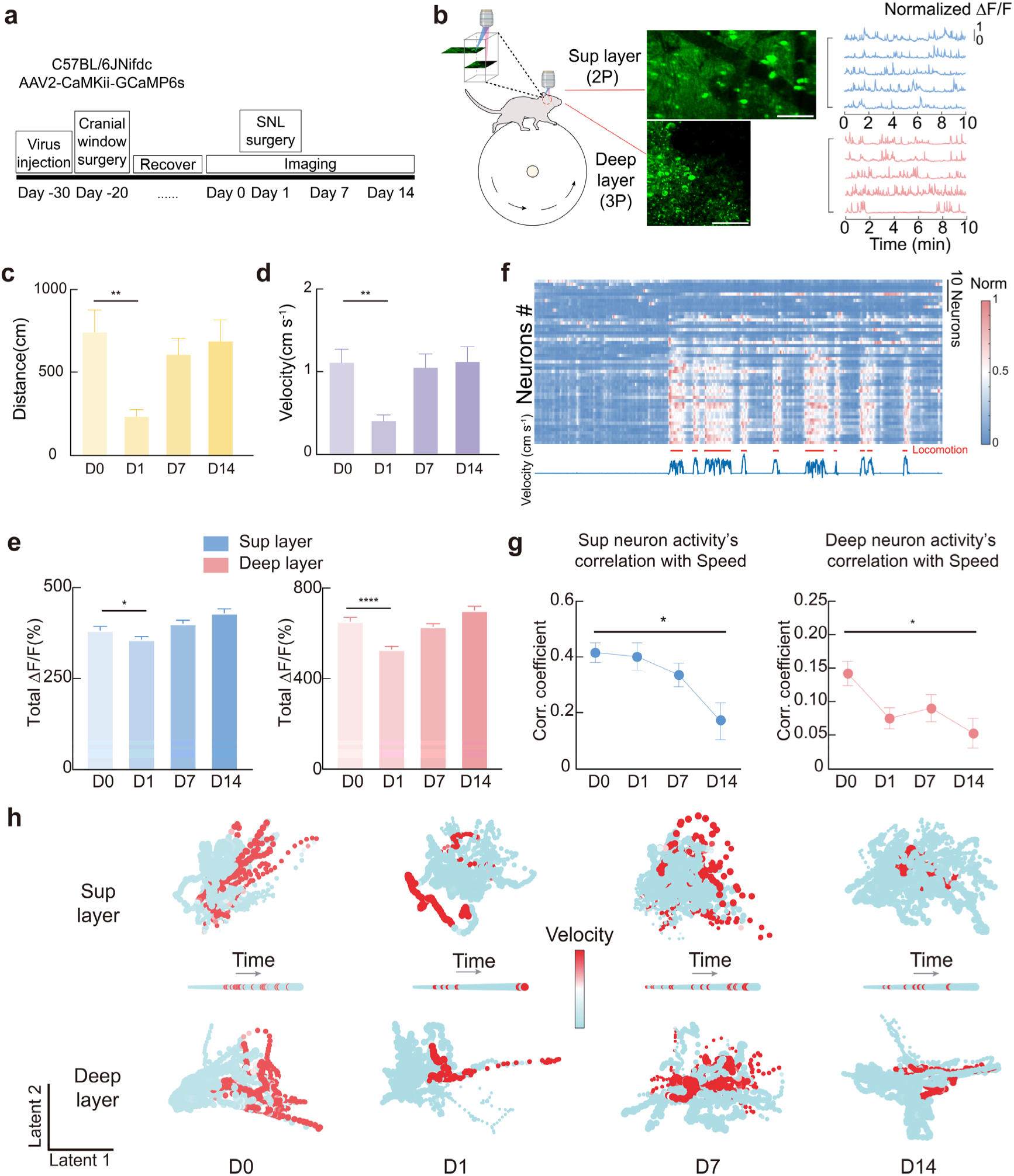
Differential responses of superficial and deep motor cortex neurons in chronic pain. a) Diagram of experimental design. b) Schematic of the mouse behavioral setup and imaging results. Left: mouse fixed on a running wheel during imaging; middle: imaging results of superficial and deep motor cortex; right: neuronal activity demonstration; scale bar: 100 μm. c) Mouse movement distance on different days. d) Mouse movement speed on different days. In panels c and d, statistical method: unpaired two-tailed t test, *n* = 10-20 FOVs (5 mice) in 4 days, *P* (c) = 0.0064, *P* (d) = 0.0031. e) Statistical analysis of overall fluorescence intensity. Left: superficial layer; right: deep layer. Statistical method: unpaired two-tailed t test, *n* (2P) = 300-400 neurons (5 mice), *n* (3P) = 600-900 neurons (5 mice) in 4 days, *P* (2P) = 0.0154, *P* (3P) <0.0001. f) Neuronal activity and behavioral traces demonstration. Top: heatmap of neuronal activity; bottom: mouse movement state and speed during imaging. g) Average response coefficient of neuronal activity to movement speed. Left: superficial layer; right: deep layer. Statistical method: Kruskal-Wallis test, *n* = 14-22 FOVs (5 mice) in 4 days, *P* (2P) = 0.0228, *P* (3P) = 0.0146. h) High-dimensional features of neurons on different days. 2D PCA dimensionality reduction of neuronal populations over time, and each point represents a time point, with smaller points indicating earlier times and larger points later times, and red points represent time points when mouse movement speed >2 cm s^-1^. Statistical results shown in c, d, and e are mean + SEM, and in g is mean ± SEM.

Locomotor analysis showed that during acute pain (D1), mean running speed decreased from 1.1 ± 0.16 cm s^-1^ to 0.39 ± 0.07 cm s^-1^ and distance traveled reduced from 748 ± 134 cm to 237 ± 44 cm (Figure 5c-d), and it confirmed SNI modeling effectively impaired locomotion. Analysis of population neural activity intensity in superficial and deep layers further showed that its trend was highly correlated with locomotor states (Figure 5e), while firing frequency exhibited no significant changes (Figure S6b), indicating chronic pain specifically modulates activity intensity rather than rhythmic properties.

Neuronal heatmaps revealed that neurons exhibited higher activity intensity during mouse locomotion (Figure 5f), suggesting more pronounced single-cell level responses to movement. Pearson correlation coefficients between single-neuron activity and running speed (Figure 5g) demonstrated that superficial layer showed peak pre-SNI responses, with significant reductions at D7 and D14, while deep layer exhibited early attenuation by D1 persisting through D14. The data results in the high-dimensional space also demonstrate similar findings (Figure 5h): at D0, both the shallow and deep regions clearly distinguish the movement state from the static state. However, at D7/D14, the state discrimination of the shallow region significantly decreases, and at D1/D7/D14, the state discrimination of the deep region also significantly decreases. FOV-based neuron classification revealed dynamic shifts in response-type proportions during pathogenesis (Figure S6c), indicating layer-divergent adaptations of movement tuning in motor cortex across acute-to-chronic pain stages. However, pain minimally affected neuronal population coordination. Population synchrony within/between cortical layers remained unchanged pre-/post-pain modeling (Figure S6d). SVM classifiers using layer-specific activity predicted locomotor states with >90% accuracy across timepoints (Figure S6e), demonstrating population-level coding dominance over single-cell functions.

In summary, simultaneously 2P and 3P imaging of superficial-layer (L2/3) and deep-layer (L5) neurons in the M1 of SNI model mice revealed that pain-induced motor dysfunction correlated with consistent trends in neuronal activity intensity. However, single-cell responses exhibited temporal divergence: superficial-layer neurons showed progressive functional decline during the chronic phase, whereas deep-layer neurons displayed impaired activity acutely. Population-level synchronization remained stable. This work delineates the functional dissociation between individual neurons and population coding in the motor cortex during locomotion, providing novel insights into the neural mechanisms by which chronic pain disrupts motor regulation.

## 3. Conclusion

Despite its demonstrated performance, several areas of the current 2PIM-3PM system merit further optimization to enhance performance and facilitate broader adoption:

First, the current configuration requires two separate laser sources for 2P and 3P excitation. Although this ensures clean spectral and temporal separation, it increases system cost and alignment complexity. A delay-line-based beam-splitting strategy is proposed (**Figure S7**) to allow dual-mode operation using a single laser, which could streamline integration and reduce maintenance overhead.

Second, while the 2PIM-3PM platform excels in functional imaging, it currently lacks optical manipulation capabilities. Fortunately, the system is compatible with holographic 2P optogenetics through dedicated beam paths, which can be integrated downstream of the scanner (**Figure S8**). Incorporating these tools will allow causal dissection of cell-type-specific and projection-specific circuit dynamics^52, 53^.

Third, the lateral translation range of the 2P imaging beam is limited by the FOV of standard objectives. Current implementations cover up to 1 mm lateral movement. Future versions could incorporate mesoscopic objectives with extended FOVs (4–5 mm), facilitating cross-structure studies, such as mPFC–hippocampus or retrosplenial– visual cortex circuits^54–59^.

Fourth, current validation focuses on two behavioral paradigms—air puff stimulation and chronic pain—chosen to demonstrate performance under transient and persistent conditions, respectively. However, the system’s utility can be further expanded by incorporating a broader range of cognitive and emotional tasks, including learning, affective state encoding, and sleep modulation. Such extensions will deepen the biological relevance of the platform and enable exploration of layer-specific computations across brain states.

In summary, 2PIM offers a compact, modular solution for high-speed, dual-depth and multi-region *in vivo* imaging, with precise control over axial and lateral positioning. By enabling simultaneous recording of distributed neural activity across cortical layers and brain regions without compromising spatiotemporal resolution, it provides a powerful platform for studying the spatiotemporal dynamics of distributed neural circuits.

## 4. Methods

### 4.1 Imaging System Configuration

#### 4.1.1 3PM Configuration

3P excitation was performed using a noncollinear optical parametric amplifier (I-OPA-TW-F, Light Conversion) operating at 1300 nm with a repetition rate of 1 MHz. Group velocity dispersion (GVD) was pre-compensated to counteract the normal dispersion introduced by downstream optical elements.

Two objectives were used interchangeably depending on the imaging requirements: a 25× water-immersion objective (NA = 1.05, Olympus) and a 16× water-immersion objective (NA = 0.8, Nikon).

Both fluorescence and third-harmonic generation (THG) signals were collected in the epi-direction through the objective lens. The signals were first reflected by a primary dichroic beam splitter (FF775-Di01-25x36, Semrock) and then separated by a secondary dichroic beam splitter (FF458-Di02-25x36, Semrock), which directed the fluorescence and THG signals to separate GaAsP photomultiplier tubes (PMTs; H7422-40, Hamamatsu). Bandpass filters (520/60 nm for fluorescence and 447/60 nm for THG, both from Semrock) were placed in front of the respective PMTs for spectral separation. The sample was mounted on a motorized stage (MP-285A, Sutter Instrument), and image acquisition was controlled via ScanImage 2021 (Vidrio Technologies) running on MATLAB (MathWorks). PMT output currents were converted to voltages using transimpedance amplifiers (C12419, Hamamatsu). A galvo-galvo scanner (6210H, Novanta) was used for beam steering. A schematic of the optical setup is shown in Figure S1.

#### 4.1.2 2PIM Integration

The 2P laser source was Coherent Chameleon Ultra2, which emits 140 fs pulses at a repetition rate of 80 MHz. The laser beam was first collimated using mirrors and then passed through an EOM and a PBS (PBS632, Lbtek) for intensity modulation. It then entered the RF submodule, which controlled axial focus. Upon exiting the RF submodule, the beam was relayed through a lens pair to the RLS submodule, responsible for XY position adjustment. Finally, the beam was coupled into the main 3P optical path via a DM (DMR-1180LP, Lbtek), allowing seamless 2P/3P integration.

#### 4.1.3 TDM Submodule

The TDM submodule utilized a 1 MHz EOM (Fujian Castech Optics, China) with a BPS-214BD-920 crystal and a PCDH-S2-1000-E-N amplifier. The optical path was configured as follows: the laser beam emitted from the laser source first passed through a half-wave plate (HWP, WPH05M-915, Thorlabs) and a PBS to adjust the polarization state (not shown in figure), then entered the EOM perpendicularly. A second PBS was used downstream to separate the modulated laser beam.

The modulation signal for the EOM was generated by an AWG (SDG 1062X, Siglent), synchronized to the clock signal of the 3P laser. The AWG output was set to 1 MHz frequency with a pulse width of 144 ns, amplitude of 4 V (high) and 0 V (low), and a delay of 820 ns. This signal was fed into the EOM amplifier to achieve precise modulation.

#### 4.1.4 RF Submodule

The RF submodule primarily consisted of a PBS, a QWP (WPQ05M-915, Thorlabs), a doublet lens (*f* = 19mm, AC127-019-B, Thorlabs), a mirror, and a VC (XRV97, Akribis) refer to the previous design^31, 60^. The modulated femtosecond laser beam first passed through the PBS and QWP, then was focused onto the mirror by the doublet lens. The mirror was mounted on the VC, enabling fast axial scanning along the Z-axis. After reflection, the laser beam retraced the same optical path through the doublet lens and QWP and was redirected by the PBS into the downstream optical path toward the microscope. The VC provided μm-scale precision in Z positioning, ensuring precise axial positioning of the 2P imaging plane.

#### 4.1.5 RLS Submodule

The RLS submodule consisted of a pair of relay lenses (*f* = 100 mm, AC254-100-B, Thorlabs; *f* = 150 mm, AC254-150-B, Thorlabs), a fixed mirror (PF10-03-G01, Thorlabs), and a motorized mirror mount (Picomotor 8816-6, Newport). The laser beam, reflected from the RF submodule, was focused onto the motorized mirror, which was conjugated to the galvanometer scanning plane via the relay lens pair. This configuration allowed remote adjustment of the imaging position in the X-Y plane. The two-axis knobs on the motorized mirror mount were individually controlled by a custom-built pulse generator that varied the direction and amplitude of the output pulses to adjust the mirror angles.

### 4.2 Optical Characterization and System Validation

#### 4.2.1 Zemax Optical Simulation

To characterize the optical performance and ensure accurate alignment of the 2PIM integrated into the 3PM system, we conducted comprehensive Zemax optical simulations. All components except the objective lens and galvanometric scanners were imported using Thorlabs’ optical libraries. In the RF submodule, the system comprising the PBS, QWP, and a pair of 19-mm doublet lenses was simplified into a relay system. Z-Axial focus shifting was achieved by adjusting the position of the first lens, thus varying the spacing between the lens pair.

For scanning simulation, a pair of mirrors was placed at the galvanometer position in the beam path, corresponding to the galvo-X and galvo-Y scanners. Tilt angles ranging from -2.5° to 2.5° were applied to the mirrors using the Multi-Configuration Editor (MCE) to simulate bidirectional scanning. To simplify the analysis, ideal lenses were used in the simulation.

For RF submodule simulation, the position of the doublet lens in the RF submodule was adjusted along the beam path, which shifted the Z-axial focal plane at the sample side after the objective. This focal plane was defined as the 2P imaging plane. The axial (Z-axis) offset between this focal plane and that of the same objective lens used in the 3PM represents the depth separation between the two imaging modalities. The scanning range under the ± 2.5° mirror tilt determined the size and shape of the resulting FOVs. For RLS submodule simulation, Adjusting the tilt angle of the conjugate mirror enables lateral displacement of the FOV on the image plane, thereby mimicking the effect of a lateral translation submodule.

#### 4.2.2 Laser Signal and Fluorescence Characterization

To measure the fluorescence signal excited by a single pixel, a 100 μM sodium fluorescein solution was prepared. Under weak laser illumination, the solution was imaged, and the output from the amplifier was connected to an oscilloscope (SDS 1202X-E, Siglent) for data acquisition. The recorded signals were subsequently analyzed using MATLAB. For validation of the fluorescent imaging, conventional images were also acquired from the same solution.

To characterize the 3P, 2P, and modulated 2P laser signals, a temporary optical setup was assembled. The laser beam was first attenuated using a neutral density filter (ND, NE30A, OD = 3, Thorlabs), then focused by a lens onto a photodetector (PD, DET08CFC, Thorlabs). The PD output was recorded by the oscilloscope, and the data were analyzed in MATLAB.

#### 4.2.3 Temporal Gating and Signal Separation

Fluorescence signals from different imaging depths are collected by a PMT after passing through the detection optical path. The resulting photocurrent is amplified using a 1 MHz amplifier and then transmitted to a data acquisition card (DAQ, MBFscience). The analog signals acquired by the data acquisition card are temporally separated using ScanImage software based on defined time windows: signals acquired 370-500 ns after laser excitation are assigned to 3P imaging, while those acquired 1,000-1,100 ns post-excitation are attributed to 2P imaging. This temporal gating effectively separates the 2P and 3P signals, allowing the generation of corresponding final images.

#### 4.2.4 Imaging Procedure

For conventional time-series imaging, 3PM was first used to locate deep regions of interest, such as the dorsal hippocampal CA1 at 1,000 μm or cortical layer at 700 μm depth. We then adjusted the 2PM position independently in both axial and lateral dimensions. The imaging depth of 2PM was first tuned by adjusting the VC in the RF submodule, until the desired superficial layer was reached. Then, the lateral position was adjusted by manipulating the motorized mirror in the RLS submodule, allowing precise targeting of the region of interest within the superficial plane.

During the alignment process, laser power was carefully controlled. The 3PM focal power was maintained below 2 mW (corresponding to a pulse energy < 2 nJ), and the total laser power was kept below 100 mW. For 2PM imaging, the laser power typically ranged from 2-20 mW, with total power remaining under 100 mW. Imaging was performed at 512 × 512 pixels per frame with a frame rate of 3.6 Hz. Single imaging sessions typically lasted 5-30 minutes; however, even 1-hour sessions were well-tolerated, with no observable tissue damage as confirmed by histological analysis. For long-term imaging, ultrasound gel was used as the immersion medium instead of water to prevent evaporation or liquid loss.

For volumetric imaging, the Z-stack function in ScanImage was employed following time-series acquisition to collect volumetric image, using a step size of 2 μm.

### 4.3 Animal Procedures

#### 4.3.1 Animals

All animal experiments were performed according to ethical compliance approved by the Institutional Animal Care and Use Committee of the Department of Laboratory Animal Science at Fudan University (2022JS-ITBR-013).

Wild-type C57BL/6JNifdc mice (male, 6-8 weeks old, from Charles River) were used for experiments (Section 2.2, 2.3 and 2.5). CaMKii-Cre (#C001015, from Cyagen Biosciences) and Ai162 mice (#031562, from Jackson Labs) were in-house bred, and heterozygous CaMKii-Cre::Ai162 mice were used for experiments (Section 2.4). Animals were housed under a 12-hour light/dark cycle with ad libitum access to food and water. All procedures were conducted by protocols approved by the Institutional Animal Care and Use Committee of the Department of Laboratory Animal Science at Fudan University.

#### 4.3.2 Viral Injection and Cranial Window Implant

Mice were anesthetized with 1-2% isoflurane and secured in a stereotactic frame (RWD 68803) on a heating pad. Erythromycin ophthalmic ointment was applied to protect the eyes. For viral injection, a pulled glass micropipette loaded in a 10 μl Hamilton syringe (Model 701N) was controlled by a microinjection pump (KDS 130, KD Scientific). Virus was delivered at 50 nl min⁻¹, and the pipette was retained for 10 minutes post-injection to prevent backflow.

For cortex-CA1 dual-region imaging, 200 nl of pAAV2/9-CaMKIIα-GCaMP6s (titer: ∼1.55 × 10¹³ viral genome copies (vg) ml⁻¹, Obiosh) was injected into the cortex (AP -1.8 mm, ML +1.5 mm, DV -0.4 mm) and hippocampal CA1 (DV -1.1 mm). For motor cortex imaging, the same virus was injected either into superficial (AP -1.0 mm, ML +0.9 mm, DV -0.3 mm) or deep (DV -0.7 mm) cortical layers.

For the mice that have been injected with the virus for 21 days, to implant a chronic cranial window, a 4 mm circular craniotomy was made using a high-speed dental drill, leaving a thin skull layer intact. After gently detaching the bone flap with forceps under saline, the dura was preserved. Bleeding was controlled with absorbable hemostatic agent. A custom 4 mm coverslip (150 μm thick) was placed over the exposed brain and sealed with tissue adhesive (3M Vetbond). A custom titanium headplate was glued to the skull using +cyanoacrylate adhesive (ergo, 5800).

#### 4.3.3 SNI Model

The SNI model was performed in adult C57BL/6 mice (8-12 weeks old, the craniotomy operation has been completed) under 2% isoflurane anesthesia. After hair removal and disinfection with 75% ethanol, a 1-cm longitudinal incision was made in the lateral thigh under sterile conditions. The biceps femoris muscle was bluntly dissected to expose the sciatic trifurcation, identifying the tibial, common peroneal, and sural nerves. Using microsurgical forceps and 20× magnification, the tibial and common peroneal nerves were doubly ligated with 8-0 non-absorbable silk sutures (2 mm apart), followed by transection and removal of the 2-mm nerve segment distal to the proximal ligature to prevent axonal regeneration. The sural nerve was meticulously preserved without mechanical manipulation.

Mechanical withdrawal thresholds were then measured using a series of calibrated von Frey filaments. Specifically, each filament was applied perpendicularly to the plantar surface of the hind paw or fore paw with sufficient force to bend it. The mechanical response threshold was defined as the minimal force filament that elicited a brisk paw withdrawal, flinching, or licking behavior. If no positive pain response occurred, a filament with a greater force was used. The measurement was repeated five times to obtain an average threshold.

#### 4.3.4 Behavioral Paradigm

We developed a head-fixed virtual reality (VR) system based on previously published designs^39, 61, 62^. The setup includes a custom-built running wheel (8 cm wide, 14 cm diameter), mounted on a 6 mm-diameter, 30 cm-long stainless rod. This rod is supported by bearings within a custom-designed bracket and connected at one end to a rotary encoder for motion tracking. The entire assembly is placed on an elevation stand (Lbtek LJ1209). During experiments, head-fixed mice are free to run forward or backward on the wheel, with their movement continuously recorded by the encoder and transmitted to an Arduino Uno microcontroller.

Virtual environments were designed in ViRMEn (Virtual Reality MATLAB Engine, 2016 release) and were presented to the mice through a 120° LED screen. Custom scripts in MATLAB can get incoming string flow from an Arduino Uno.

For air puff stimulation, an air puff system was implemented. Compressed air is delivered through a three-way solenoid valve, which by default vents to the environment. Upon receiving a control current, the valve redirects airflow to a stimulus delivery tube. The valve is gated by a custom MATLAB script, which synchronizes stimulus timing with image acquisition. Specifically, the frame clock from ScanImage is routed to the Arduino via a DAQ, allowing precise temporal alignment between behavioral data and imaging frames. The Arduino integrates encoder and timing signals to trigger air puff delivery at designated time points. For both visual and air puff stimuli, Custom scripts in ViRMEn were used to interface the animal’s behaviour with the VR: incoming TTL pulses from a rotary encoder recorded animal position to update the screen and use the MATLAB clock to determine when to give a blow stimulus or change environment. Illumination from the LED screen can pass through the objective lens and enter the PMT, thereby contaminating the imaging signal by introducing background noise. As a precaution during visual stimulation experiments, we encapsulated the objective lens with commercially available black modeling clay to block interference from the LED illumination on the imaging.

To acclimate mice to the imaging environment and running wheel, head-fixed mice were allowed to familiarize themselves with the setup for 30 minutes daily for 5 consecutive days, during which they gradually learned to locomote on the wheel.

#### 4.3.5 Histology and Immunohistochemistry

By the end of each experiment, mice were transcardially perfused with ice-cold saline, followed by 4% paraformaldehyde (PFA). Brain was swiftly removed and postfixed in 4% PFA at 4 ℃ overnight. After cryoprotection in 30% sucrose for at least 24 h, the brain was next embedded in optimal cutting temperature compound (OCT, Sakura Finetek) and sectioned at 30 μm thickness on a cryostat (Leica Microsystems). For immunochemistry staining, brain sections were washed with PBS (in this section, PBS refers to phosphate buffer saline) for 5 minutes and then blocked in blocking solution, containing 5% normal donkey serum (Jackson) and 3% Triton X-100 (Sigma) in PBS, for 30 minutes at room temperature (RT). Sections were incubated with the primary antibody at 4 ℃ overnight. After rinsing in PBS (10 minutes ×3), sections were incubated in DAPI (1:1000) and secondary antibody at RT for 2 h. Following staining, slices were washed in PBS (10 minutes ×3) before being mounted on slides and coverslipped with 70% glycerol in PBS. To check the expression of GCaMP6s, primary rabbit anti-GFP antibody (1:500) (Invitrogen A-11122) and secondary Alexa Fluor 488-conjugated donkey anti-rabbit (1:500) (Jackson 711-545-152) were used. To check the expression of Hsp70, primary mouse anti-Hsp70 antibody (1:400) (Enzo Life Science ADI-SPA-810-D) and secondary Alexa Fluor 594-conjugated donkey anti-mouse (1:500) (Jackson 715-585-151) were used.

### 4.4 Data Processing and Analysis

#### 4.4.1 Preprocessing

All acquired imaging datasets were processed using a custom-developed preprocessing pipeline. First, PMT ripple noise with gray values below 700 was automatically removed, followed by rigid image registration to correct motion artifacts. Subsequently, DeepCAD was applied for deep-learning-based image denoising^63^. Cell detection and calcium signal extraction were then performed using Suite2p^64^ (https://github.com/MouseLand/suite2p).

For single-neuron activity traces, the baseline fluorescence (*F_0_*) was calculated as the average signal within the 10-30% lowest percentile of the trace. The relative fluorescence change (Δ*F/F*) was computed using the formula:

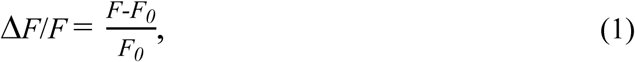

where *F* represents the raw fluorescence signal, and *F_0_* is the calculated baseline.

Based on Δ*F/F*, referring to previous studies^65^, we defined a peak exceeding the mean + two standard deviations as a calcium event. Then the firing frequency is calculated by dividing the number of events by the corresponding time. And the activity intensity of a period of time is the sum of the Δ*F*/*F* values of this period.

4.4.2 *Photon Count Estimation*

To estimate the number of fluorescence photons collected during imaging, we first calibrated the detection system such that a single detected photon corresponded to an average grayscale value of 387. This conversion factor was derived from single-photon response calibration using fluorescein. At a given time point t, the total number of photons detected within a region of interest (ROI) was estimated as: Photons = (ROI Area × ROI Mean) / 387, where ROI Area is the total number of pixels within the ROI, and ROI Mean is the average grayscale intensity of those pixels.

#### 4.4.3 d’ Signal Quality Assessment

We calculated the *d’* values for 2P, 3P, and background-subtracted 2P signals based on the method described by Mark J. Schnitzer^34^. The calculation follows the equation:

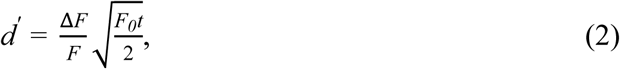

where Δ*F/F* is the amplitude of the optical transient (assumed to be 0.3 for GCaMP6s), *t* is the duration of the calcium fluorescence signal change induced by a single action potential (for GCaMP6s, *t* = 1.5).

#### 4.4.4 BS-2P Processing

we assumed the background fluorescence to be a uniform DC component across the entire image field. By contrast between the neuron and the background adjacent to the neuron in a single frame image, a fixed grayscale offset of 100 was subtracted from raw 2P images to generate BS-2P data. Subsequent photon counts, *d′*, and Δ*F/F* calculations were performed on BS-2P group.

#### 4.4.5 Photobleaching Quantification

To evaluate fluorescence decay during simultaneous 2P/3P long-term imaging. Bleaching curves were computed per neuron using a sliding window approach (window: 200 frames ≈ 1 minute) and the 30th percentile within each window was selected to minimize outlier effects as the value of this window. Group-level curves represent mean *±* SEM across neurons.

#### 4.4.6 Heat-Induced Tissue Damage Assessment

To quantitatively assess heat-induced tissue damage, based on the method described by Thornton^23^, gray values were measured in three distinct regions: the imaging area, a contralateral control area in the opposite hemisphere, and regions exhibiting surgical injury near the cranial window. Statistical method is unpaired two-tailed t-tests to evaluate significance.

#### 4.4.7 Motion State Extraction

Due to the mismatch between the sampling rate of the rotary encoder and the imaging frame rate, encoder timestamps were first downsampled in MATLAB to match the imaging frame clock. Displacement data were then aligned using these downsampled timestamps to generate a motion trace synchronized with the imaging frames. Instantaneous motion velocity was subsequently calculated from the downsampled displacement data.

#### 4.4.8 Correlation-Based Analysis

To assess neuronal responses to motion and air puff stimuli, Pearson’s correlation coefficients were computed between individual neuronal traces and either motion velocity or the air puff timing sequence.

Based on Pearson’s correlation *r*, we did different analyses. 1) Responsive neuron selection: neurons stratified by stimulus-response Pearson *r*; 2) Population response intensity: mean *r* across neurons; 3) Statistical differences in different layers: Distributions of *r* in superficial vs. deep layer, and statistical method is Kolmogorov-Smirnov test; 4) Heatmap: Neurons sorted by *r* values and then plotted. 5) Classification of neurons: Neurons were classified as: positive response: *r* > 0.3; negative response: *r* < -0.3; no response: -0.3 ≤ *r* ≤ 0.3.

#### 4.4.9 Population Synchrony Analysis

To assess synchrony within and across brain regions or layers, pairwise *Pearson’s correlation* coefficients were calculated: Within-layer: Among N neurons in layer X (N×(N-1)/2 pairs). Cross-layer: between N layer X and M layer Y neurons (N×M pairs). Statistical method is Kruskal-Wallis test.

#### 4.4.10 Temporal Sequence Analysis of Air Puff Response

To analyze timing differences in responses between superficial and deep layers neurons following air puff stimulation. 1) Signal averaging: for each FOV, activity traces of positive responders in each layer were averaged. 2) Select time window: a 30 s window (10 s pre-, 10 s during, 10 s post-stimulus) was analyzed. 3) Normalization: averaged traces were normalized to the range [0,1]. 4) Calculate rise time: defined as the time first to reach 0.5 after stimulus onset. 4) Calculate decay time: defined as the time to decay from 1 to 0.5. 6)Temporal difference: the Δ (rise time) and Δ (decay time) between layers. The statistical method is the Kruskal-Wallis test.

To evaluate response stability between the with air puff and the without air puff conditions, the coefficient of variation (*CV*) was computed for each dataset using the formula:

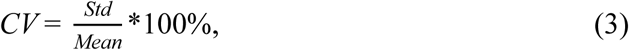

#### 4.4.11 Latency Analysis

Neuronal events were identified following the method of Nakul Yadav^13^. An event was defined as a period where Δ*F/F* exceeded 3*σ* for at least two consecutive frames. Inactive neurons were excluded. The onset times of active neurons were plotted as cumulative distribution functions, and group differences were assessed using the Kolmogorov-Smirnov test.

#### 4.4.12 Dimensionality Reduction Using PCA

To visualize population-level neuronal dynamics in a reduced feature space, PCA was applied using MATLAB’s built-in function. For sensory cortex data, activity traces from superficial and deep neuronal populations were analyzed along the time dimension. The first two principal components (PC1 and PC2) were extracted and plotted in 2D, with timepoints color-coded by the air puff stimulus vector or by time on a continuous scale.

Similarly, for motor cortex recordings following pain model induction, PCA was used to project time-resolved population activity from superficial and deep layers within individual FOVs onto a 2D space. Here, timepoints were color-coded based on locomotion speed, allowing for visualization of behaviorally relevant neural state trajectories across imaging sessions.

#### 4.4.13 Decoding using an SVM

To decode locomotion states from cortical population activity, a linear-kernel SVM classifier was trained on calcium signals at each day. For both superficial and deep layers within a given FOV: 1) Data structuring: Neuronal activity matrix: *N* neurons × *T* timepoints; Locomotion label: Binary vector *V* (movement: velocity > 2 cm s^-1^ = 1; rest: velocity ≤ 2 cm s^-1^ = 0); Under sampling applied to balance movement/rest durations → *V_1_* (1×*T_1_*). 2) Model workflow (exemplified by D0 superficial layer): Stratified split: 70% training / 30% testing (*n* FOVs); SVM trained on training set; Testing accuracy = mean prediction correctness.

### 4.5 Statistical Analysis and Quantification

Data are presented as mean ± SEM or mean + SEM. Statistical analyses were performed using MATLAB (MathWorks, R2022b) and SPSS (IBM, v20). Significance thresholds were set at \**P* < 0.05 and \*\*\*\**P* < 0.0001. Detailed statistical information, including the exact value and definition of *n*, is provided in the corresponding figure legends.

## Acknowledgement

This work was supported by the National Natural Science Foundation of China (T2222006, 32471142, 32200919), Fudan University (yg2023-04, FudanX24AI048), and the Shanghai Municipal Education Commission (2024RGZNB03).

## Author Contributions

B.L. supervised the project and co-designed the study with Y.G.Z. F.X provided the Zemax software (Zemax OpticStudio 2022) for optical simulation. Y.G.Z. and C.H.K. performed the optical. Y.G.Z. mechanical design, built the imaging system, and conducted the imaging experiments. J.X.S. developed the data preprocessing software. Y.G.Z. and S.L. analyzed the data. Y.G.Z., X.Y.G., and Y.F.Z. performed the animal surgeries. Y.G.Z., B.L., and X.Y.G. wrote the manuscript.

## Competing interests

The authors declare no competing interests.

## Data Availability Statement

The data that support the findings of this study are available from the corresponding author upon reasonable request.

All the codes used for data processing will be uploaded to Github (MultiRegionImaging/2PIM-3PM-Imaging).

**Correspondence** and requests for materials should be addressed to B.L.

## Notes

Funding: This work was supported by the National Natural Science Foundation of China (T2222006, 32471142, 32200919), Fudan University (yg2023-04, FudanX24AI048), and the Shanghai Municipal Education Commission (2024RGZNB03).

### Competing Interest Statement

The authors have declared no competing interest.

