## Supplementary Files for "A Compact Two-Photon Module for Simultaneous Dual-Depth, Multi-Region *in vivo* Brain Imaging"

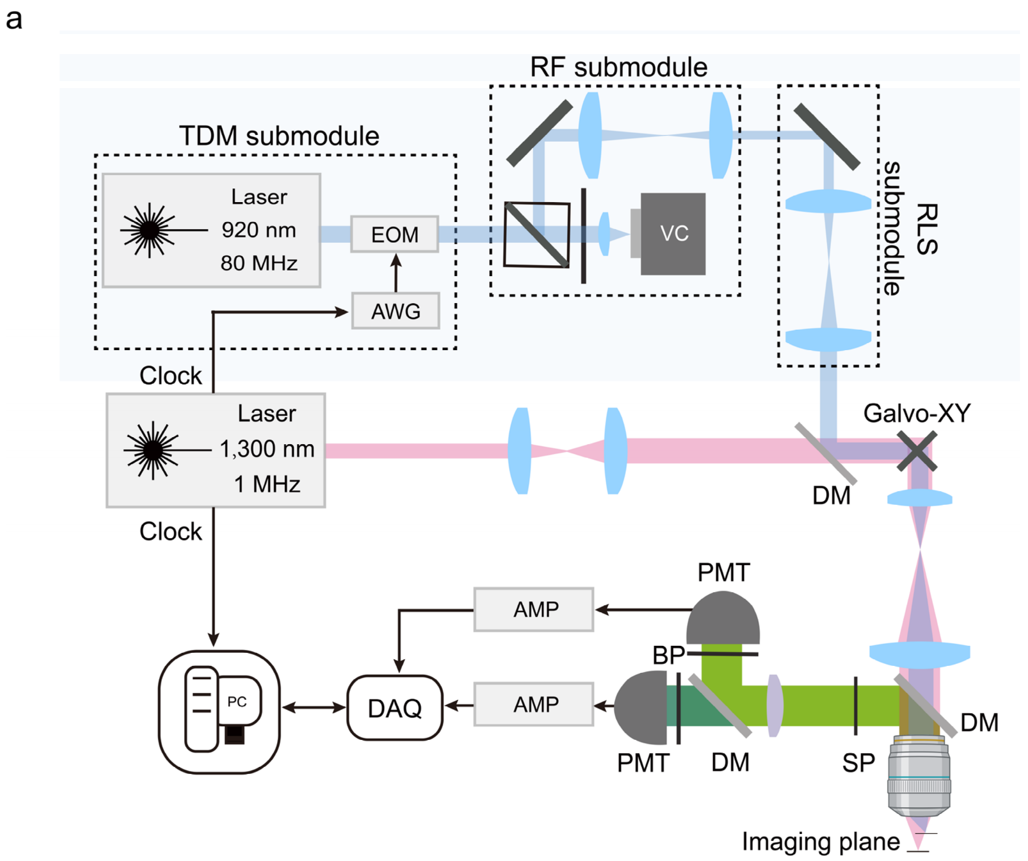


**Figure S1.** Overview and schematic of the compact two-photon imaging module integrated into a 3PM. a) Schematic of the compact 2P imaging module integrated into 3PM. EOM: electro-optic modulator. AWG: arbitrary waveform generator; VC: voice coil; DM: dichroic mirror; DAQ: data acquisition; AMP: amplifier; PMT: photomultiplier tube; BP: band pass; SP: short pass. The conventional 3P system includes standard excitation light paths and a fluorescence collection system, while the compact 2PIM comprises TDM, RF, and RLS submodules. The red path indicates 3P excitation, and the blue path indicates 2P excitation.

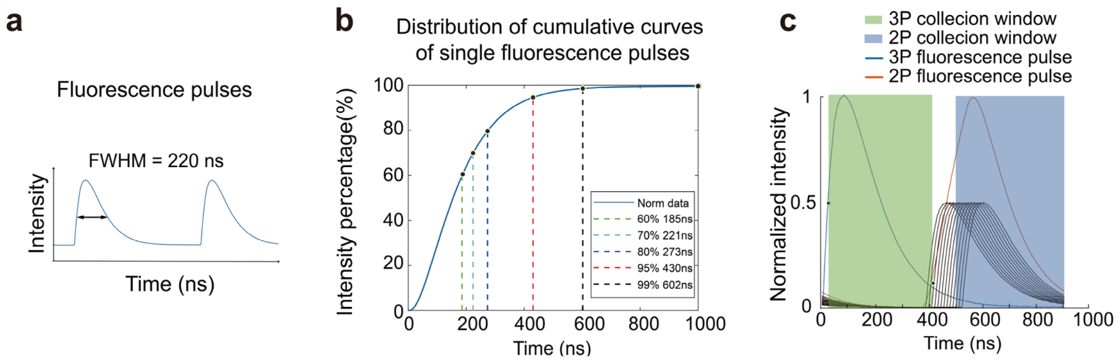


**Figure S2.** Fluorescence pulses collected from the existing three-photon system and analysis results. a) Single-pulse fluorescence signal. A single-pulse fluorescence signal measured in a fluorescent solution using the existing system, with a full width at half maximum (FWHM) of ~220 ns. b) Integrated fluorescence intensity of a single pulse. c) Simulated calculations for optimal excitation and collection ratios. The blue curve represents a single three-photon fluorescence pulse, the black curve shows 13 independent 2P fluorescence pulses, and the orange curve indicates the combined fluorescence pulse curve of 13 2P pulses, along with corresponding collection windows. Calculated 2P-Noise/3P fluorescence collection = 2.34%; 3P-Noise/2P fluorescence collection = 2.41%. Parameters will be optimized in subsequent experiments to adjust for optimal imaging.

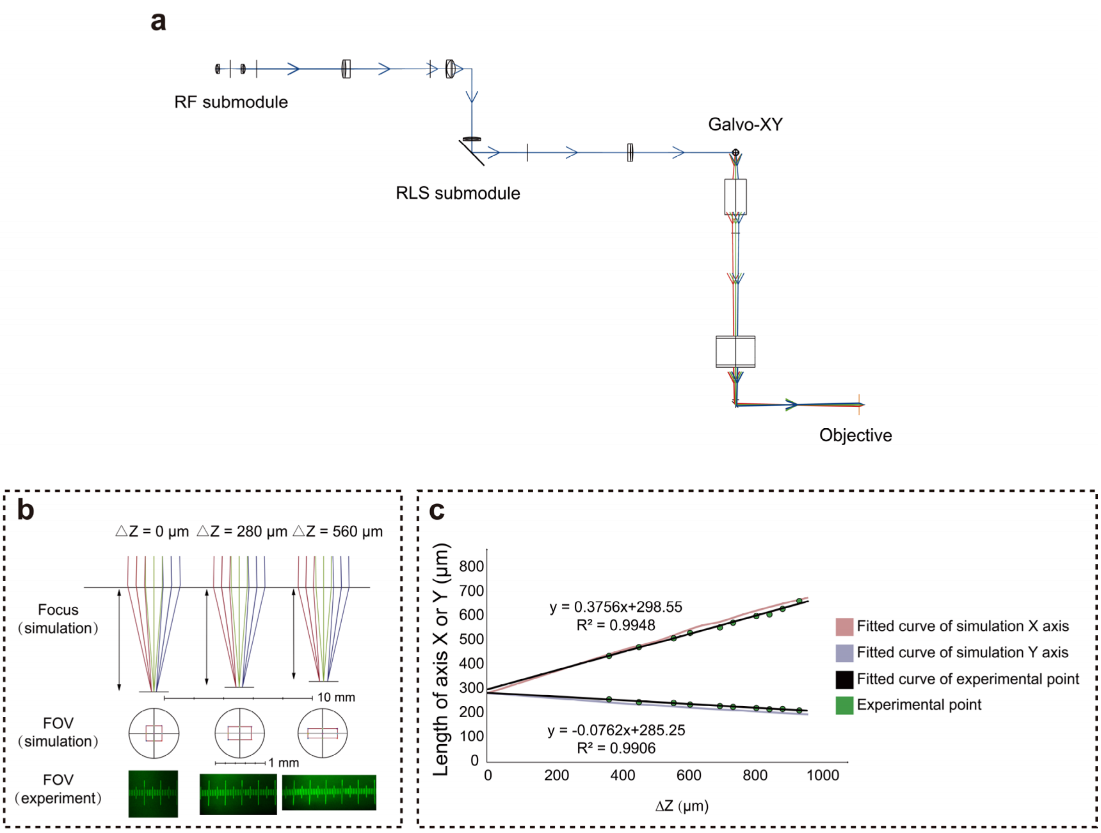


**Figure S3.** Zemax optical simulation design. a) Overall excitation optical path simulation. Zemax optical path design of the three-photon path after the RF submodule, RLS submodule, and DM. Blue indicates the 2P path, and multi-colored beams after the Galvo represent different scanning angles. b) Zemax simulation. Top: beam status and distance from the 3P imaging plane when the simulated 2P FOV moves to different depths along the Z-axis; middle: shapes of the 2P FOV at different depths; bottom: ruler image from actual imaging to show deformation effects. c) Fitted data. Comparison of fitting curves between simulated deformation data and actual imaging deformation data: red curve = X-direction simulated data fitting curve; blue curve = Y-direction simulated data fitting curve; green points = actual X/Y dimensions from imaging; black curve = fitting curve for green points, with equations above and below the black curve.

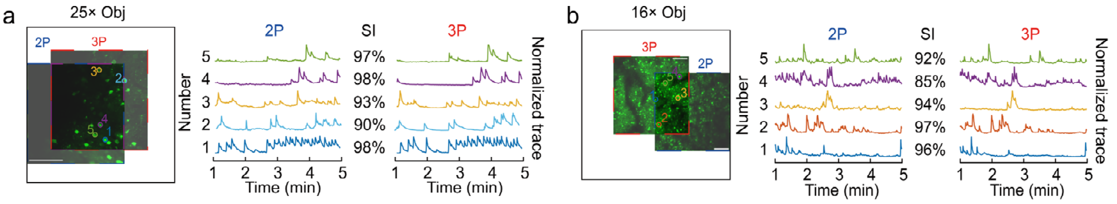


**Figure S4.** Demonstration of lateral movement capabilities of the two-photon imaging field of view. a) Lateral movement demonstration of 25× objective. Left, demonstration of 2P field moving left; Middle and Right, neuronal signal extraction from the overlapping region imaged by both 2P and 3P after movement. SI: similarity index. b) Lateral movement demonstration of Nikon 16× objective. Left, demonstration of 2P field moving right; middle and right, same as above. Scale bars: 100μm.

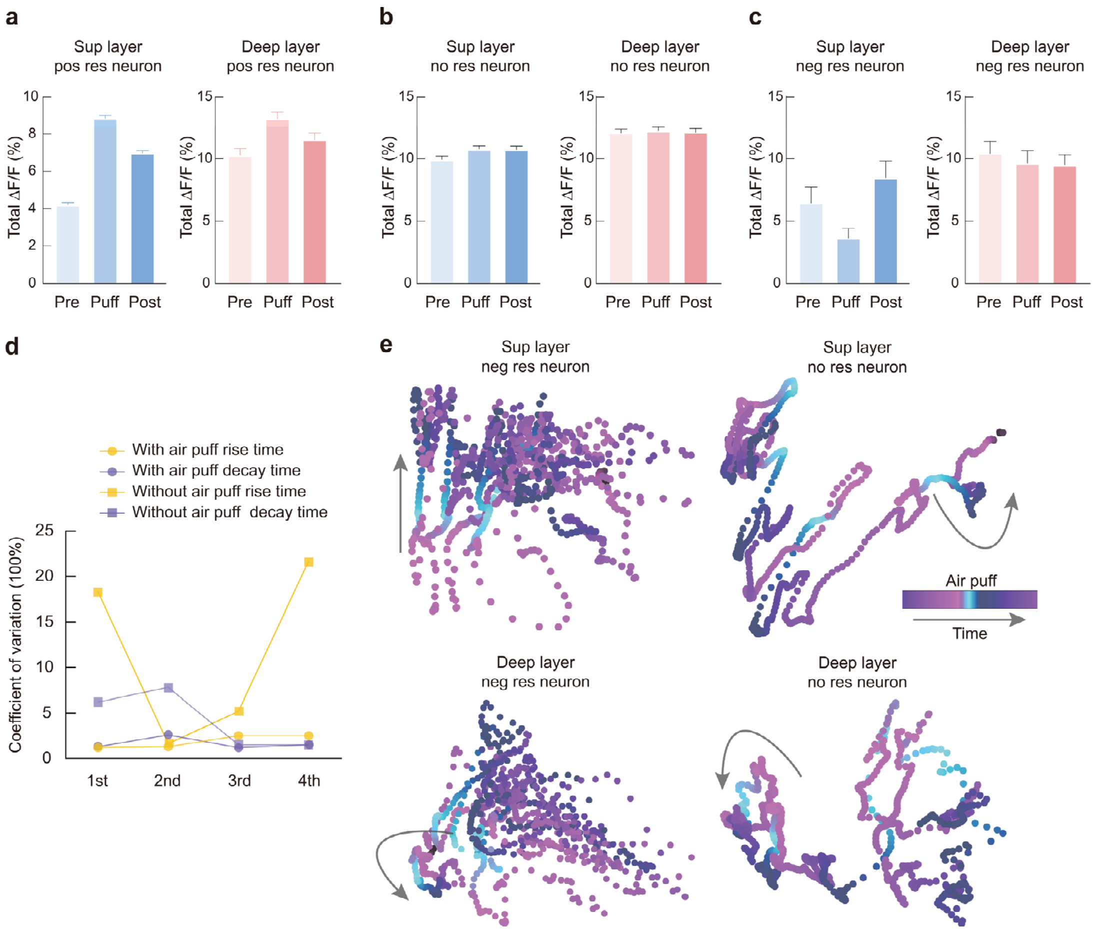


**Figure S5.** Functional differences in superficial and deep neuronal activities in the somatosensory cortex induced by air-puff stimulation using 2PIM. a) Total activity intensity of positively responsive neurons during 10 s before, 10 s during, and 10 s after stimulation; blue: superficial layer, orange: deep layer. b) Results for non-responsive neurons. c) Results for negatively responsive neurons. d) Coefficient of variation (CV) of temporal differences in rising and decaying phases of neuronal activity between superficial and deep layers during air-puff stimulation and non-stimulus time. e) High-dimensional space characteristics of non-responsive and negatively responsive neurons: Top left, 2D PCA dimensionality reduction of superficial negative response neurons over time (blue = 4 stimulus periods); bottom-left, deep negative response neurons; top-right, superficial non response neurons; bottom-right, deep non response neurons.

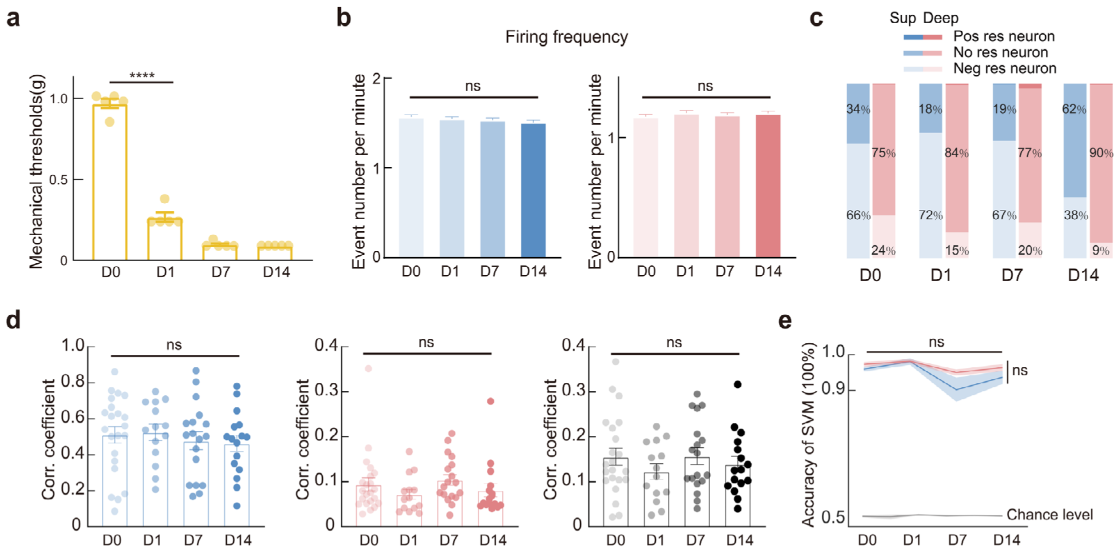


**Figure S6.** Differential responses of superficial and deep motor cortex neurons in chronic pain. a) Quantitative assessment of SNI modeling efficacy. statistical method: two-tailed paired t-test, $n = 5$ mice, $P < 0.0001$. b) Firing frequency of motor cortex neurons on different days. Left: superficial cortex; right: deep cortex. statistical method: Kruskal-Wallis test; $n = 14$ to 22 FOVs (5 mice); $P$ (2P) = 0.6739, $P$ (3P) = 0.1215, $P$ (2-3P) = 0.6809. c) Proportions of different neuron types in superficial and deep layers. d) Activity synchrony among neuronal populations. Left, average Pearson correlation coefficient within superficial layer neurons; middle, within deep layer; right, between superficial and deep layer neurons. statistical method: Kruskal-Wallis test; $n = 14$ to 22 FOVs (5 mice); $P$ (2P) = 0.6739, $P$ (3P) = 0.1215, $P$ (2-3P) = 0.6809. e) SVM prediction accuracy of movement states: Blue and orange represent prediction results from superficial and deep layer neurons, respectively; gray represents prediction results from random data. data plotted as mean ± sem. statistical method: Kruskal-Wallis test; $n = 9$ to 18 FOVs (5 mice); $P$ (2P) = 0.4244, $P$ (3P) = 0.3074. Statistical result in a, d and e is mean ± SEM.

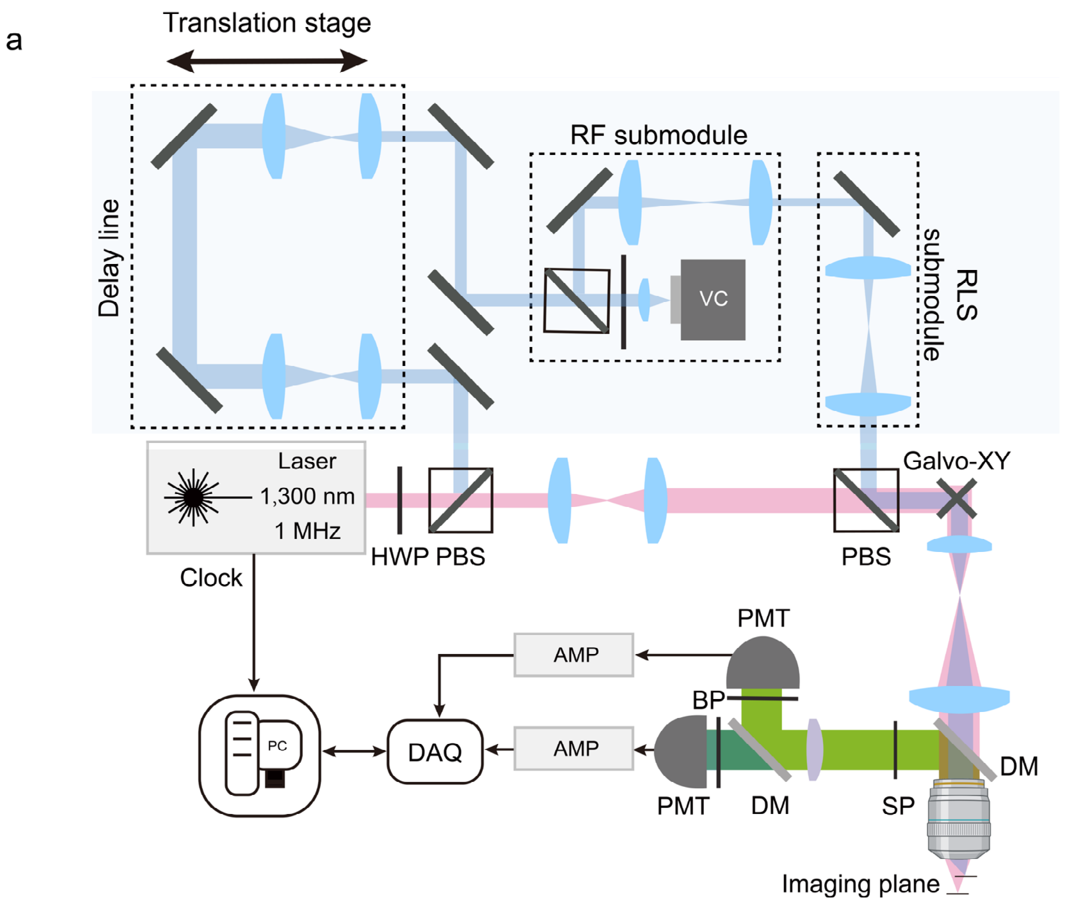


**Figure S7.** Overview and schematic of the delay-line-based beam-splitting module integrated into 3PM. a) Schematic of the delay-line-based beam-splitting module integrated into 3P. HWP: half wave plate; PBS: polarization beam splitting; DM: dichroic mirror. DAQ: data acquisition. AMP: amplifier. PMT: photomultiplier tube. BP: band pass. SP: short pass.

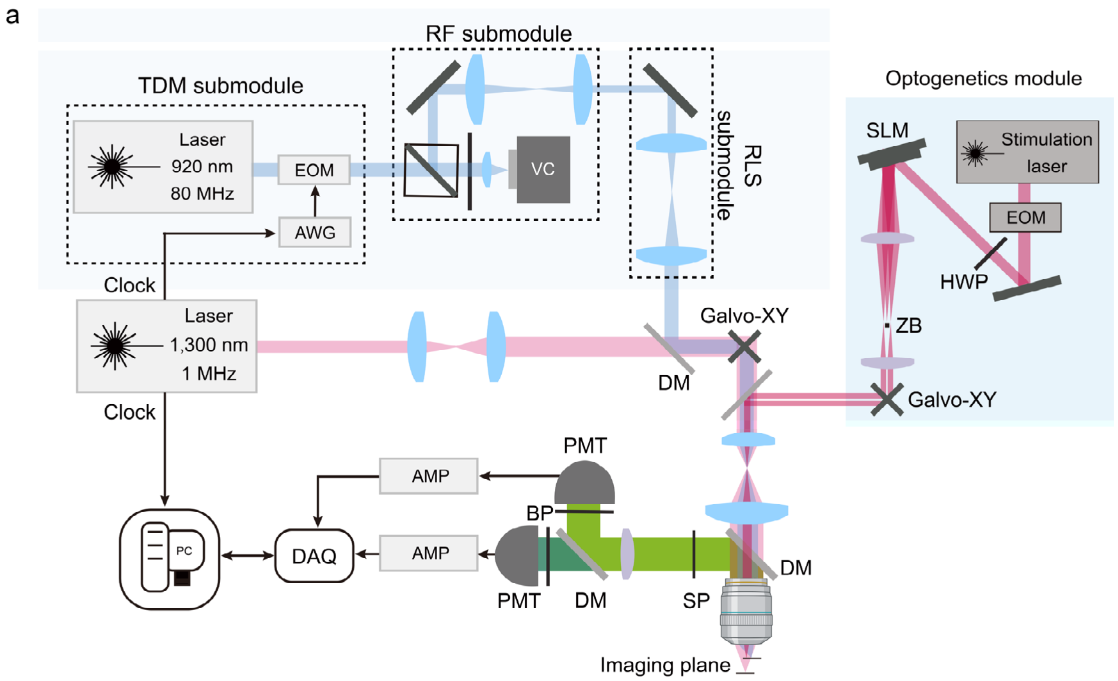


**Figure S8.** Overview and schematic of the 2PIM-3PM platform with an optogenetics module. a) Schematic of an optogenetics module integrated into 2PIM-3PM. EOM: electro-optic modulator. AWG: arbitrary waveform generator; DM: dichroic mirror; DAQ: data acquisition; AMP: amplifier. PMT: photomultiplier tube; BP: band pass. SP: short pass; HWP: half wave plate; SLM: spatial light modulator; ZB: zeroth order beam block.
